# Cholesterol plays a decisive role in tetraspanin assemblies during bilayer deformations

**DOI:** 10.1101/2021.07.29.454363

**Authors:** Marcelo Caparotta, Diego Masone

## Abstract

The tetraspanin family plays key roles in many physiological processes, such as, tumour invasion, cell motility, virus infection, cell attachment and entry. Tetraspanins function as molecular scaffolds organised in microdomains with interesting downstream cellular consequences. However, despite their relevance in human physiology, the precise mechanisms of their various functions remain elusive. In particular, the full-length CD81 tetraspanin has interesting cholesterol-related properties that modulate its activity in cells. In this work, we study the opening transition of CD81 under different conditions. We propose that such conformational change is a collaborative process enhanced by simultaneous interactions between multiple identical CD81 tetraspanins. With molecular dynamics simulations we describe the crucial role of a ternary lipid bilayer with cholesterol in CD81 conformational dynamics, observing two emergent properties: first, clusters of CD81 collectively segregate one tetraspanin while favouring one opening transition, second, cumulative cholesterol sequestering by CD81 tetraspanins inhibits large membrane deformations due to local density variations.

## 1 Introduction

Tetraspanins are a particular kind of membrane proteins expressed in animals, plants and fungi ^34,62^. Although they are the largest family of transmembrane proteins in mammals, little is known about their molecular mechanisms of action ^76^. Having a strong propensity to multimerise ^47^ they self-organize along the plasma membrane in high density clusters (tetraspanin-enriched microdomains or tetraspanin webs) ^21,33^ that play a central role in various cellular functions.

Tetraspanins are known as molecular facilitators and master organizers of the membrane architecture ^34,62^, controlling fundamental immune cell functions ^12^ and a wide range of physiological processes ^69^. They are of key importance in cell morphology, invasion, motility, fusion and signalling, in the brain, in the immune system and in tumours ^7,8,29,30,65^. Tetraspanins are involved in cell proliferation ^34,62^, pathogen uptake ^12^ and fertilization ^69^. Therefore, there is great interest in describing tetraspanins as novel drug-targets for infectious diseases and cancer ^12^.

In particular, CD81 (cluster of differentiation 81), or Tspan28^33^, is involved in cell-surface assembly of the human immunodeficiency virus (HIV), the influenza A virus and the hepatitis C virus (HCV) ^55^. Several studies have highlighted the affinity of CD81 for highly curved membranes ^23,28,75^ or its influence in membrane remodelling and deformation ^17^. Therefore, among tetraspanins, dynamic molecular characteristics of CD81 are of major interest to understand its disease-related physiological and pathological processes ^55^.

Structurally, tetraspanins contain four membrane-spanning domains connected extracellularly by a large and a small loop ^27^. Tetraspanins interact directly with lipids and other membrane proteins to control the architecture of the membrane. In particular, CD81 tetraspanin (PDB ID: 5tcx) contains a cholesterol binding pocket between its transmembrane domains which has been shown (both experimentally and computationally) ^66,76^ to influence the open/close conformational change whether cholesterol is present or not. Such opening event increases the flexibility of the external region containing γ and δ loops (res 158-174 and res 176-189 respectively), known to be important in CD81:CD81 direct interactions ^34,35,62^. In this sense, molecular dynamics has proved indispensable in both pure and applied research providing the methodology for detailed microscopic modelling on the molecular scale ^59^.

In this work, we studied CD81 assemblies as a collective phenomenon, observing their collaborative properties due to simultaneous interactions between multiple tetraspanins. First, we showed that the conformational opening transition already described for CD81 inserted in a lipid bilayer ^55,76^, takes place only partially in an isolated tetraspanin outside the bilayer. Second, we described this opening transition as favoured by assemblies of multiple CD81 tetraspanins solvated in explicit solvent, still outside the bilayer. Third, we observed that when inserted in a lipid bilayer containing cholesterol, CD81 tend to cluster and to segregate one tetraspanin. Moreover, we determined that the segregated tetraspanin shows increased openness and though higher propensity to the conformational opening transition, an emergent property not predictable by analysing an individual protein ^6,22,25^. Finally, we described the effects of CD81 tetraspanins in bending the bilayer, showing that cholesterol sequestering prevents large deformations induced by local density variations.

## 2 Results and discussion

### 2.1 CD81 open transition is a collaborative event

To observe the conformational change of CD81 from closed to open states, we have obtained the free-energy profile describing this transition in the full-length human CD81 tetraspanin (PDB ID 5tcx) ^76^ after removing cholesterol molecules (apoprotein) and solvated in TIP3 water molecules under the CHARMM36 force-field. The reaction coordinate used to explore the energy path was the 3D distance between Cα atoms in residues Phe58 and Phe126. This convenient distance definition was originally proposed by Blacklow and collaborators ^76^ to characterise, with unbiased molecular dynamics, the opening transition of a single CD81 tetraspanin inserted in a 124 palmitoyloleoyl-phosphatidylcholine (POPC) lipid bilayer, using TIP3 water molecules and the CHARMM36 force-field, as well. They showed that in 3 out of 9 trajectories the apoprotein successfully adopts an open conformation in ≈ 500ns of simulation time. This result was also repeated by Palor et al. ^55^ using a 148 POPC lipid bilayer, TIP3 water molecules and also the CHARMM36 force-field, in 500ns unbiased molecular dynamics.

Here, we have conducted umbrella sampling simulations ^60,67^ for Potential of Mean Force (PMF) calculations, spanning the Phe58:Phe126 Cα distance (*d*) from 0.5nm to 2nm to induce the opening transition of a single CD81 tetraspanin in water, without the bilayer. Molecular dynamics simulations for the opening path were biased with harmonic restrain of the form 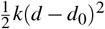, where *d* is the Phe58:Phe126 Cα distance, *k* is the force constant (in kJ/mol/nm^2^) and *d*_0_ is the equilibrium value for each umbrella sampling window. See supplementary information for technical details on umbrella sampling simulations. Panel 1a shows the free energy profile. The free energy landscape is dominated by an absolute free energy minimum with two wells at *d* ≈0.5nm and *d* ≈0.65nm. There are two other local minima at *d* ≈1nm and *d* >2nm, where feasible although metastable conformations can exist. We have measured the angle (*θ*) formed between Cα atoms of residues Val114, Val136 and Gln88 to establish effective disengagement of the extracellular large and a small loops during the opening transition (see snapshots insets in panel 1a with Cα atoms of Val114, Val136 and Gln88 residues highlighted as orange spheres).

Although conformations with distances well into open-state values were visited during umbrella sampling (*d* ≈2nm), *θ* remained in the [70,90] degrees interval (see panel 1b), indicating rather poor articulation of the Val114:Val136:Gln88 hinge and not a complete transition into the open state. Panel 1b maps all pairs of distance-angle points for each conformation adopted during all umbrella windows. Although most conformations remain in the ≈[80,100] degrees region, some states with larger angles exist (≈140 degrees). However, these conformations correspond to distances *d* ≈1.6nm in the highest region of the energy profile which belongs to transient states less likely to produce biologically representative structures.

We also performed *μ*s-length unbiased atomistic molecular dynamics of a single CD81 tetraspanin in water. Panel 1c shows all distance-angle points for each conformation visited during this simulation time. Averaged distributions for the distance and the angle are indicated in panel 1d and the averaged distance < *d* > over time is plotted in panel 1f. The combined observation of distance *d* and angle *θ* shows that the opening event does not completely occur for a single CD81 tetraspanin in water with no bilayer. Besides, after 1*μ*s of unbiased molecular dynamics the averaged distance < *d* > hardly reaches ≈0.6nm, indicating that conformations have not yet passed the second well of the absolute minimum (*d* ≈0.65nm). Panel 1e shows molecular dynamics snapshots of the initial (t=0*μ*s) and final (t=1*μ*s) states of the unbiased simulation. Consequently, we suggest that the large conformational change needed to fully open one CD81 tetraspanin might be highly facilitated by protein-lipid interactions, in agreement to others observations ^55,76^.

To evaluate whether this opening event could also driven by the interactions with other CD81 tetraspanins, we have performed additional unbiased atomistic simulations including more CD81 tetraspanins. We have run unbiased molecular dynamics for two and four CD81 equidistant replicas solvated in explicit solvent and initially disposed in line (see figure 2). Again, each structure corresponded to the full-length human CD81 tetraspanin (PDB ID 5tcx) ^76^ in its closed conformation, after removing cholesterol molecules (apoproteins).

Panel 2a plots the Phe58:Phe126 averaged distance of a system including 2 identical CD81 tetraspanins initially separated. Importantly, one of the two tetraspanins reaches distances *d* ≈0.75nm (black curve) in a relative short simulation time (more than an order of magnitude less, in comparison to figure 1f). Panel 2b maps averaged distributions for *d* and *θ* of all conformations visited during these unbiased simulations, see supplementary information for 2D scattered maps. Panels 2c depicts final (t=100ns) molecular dynamics snapshots highlighting the plasticity of γ and δ loops (res 158-174 and res 176-189 respectively) during this first-stage opening (see supplementary information for the initial configuration). Overall, the presence of another protein facilitates an initial transition of one of them with much more easiness than the singletetraspanin system. Undoubtedly, the free energy landscape may be also modified by the presence of second tetraspanin, simplifying an initial opening transition.

**Fig. 1.**
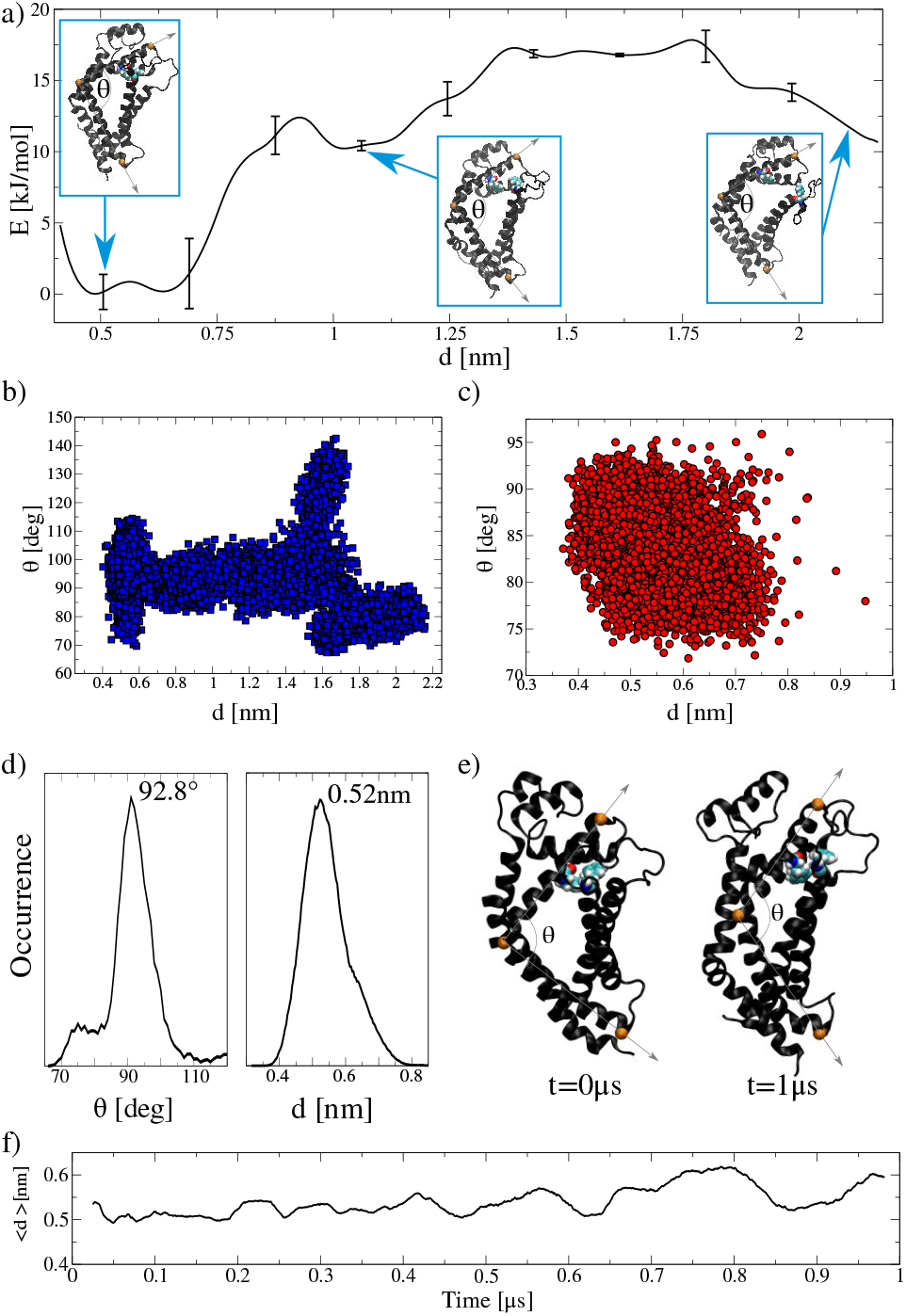
Open transition in a single CD81 tetraspanin. **a)** Free energy profile using Phe58:Phe126 Cα distance as reaction coordinate (*d*). Molecular dynamics snapshots depict the transition steps. Angle *θ* is measured between Cα atoms of residues Val114, Val136 and Gln88 (orange spheres). Error bars are standard errors calculated by individually splitting the profile in independent blocks. See supplementary information for PMF convergence details. **b)** Distance-angle 2D scattered map for all umbrella windows used to recover the free energy profile in panel a). **c)** Distance-angle 2D scattered map for 1*μ*s unbiased molecular dynamics of a single CD81 in water. **d)** Averaged distribution of the angle and the distance along the 1*μ*s unbiased simulations. **e)** Molecular dynamics snapshots show unbiased initial (t=0*μ*s) and final (t=1*μ*s) states. CD81 is represented in black ribbons, with Phe58 and Phe126 highlighted in vdW representations following the Corey-Pauling-Koltun (CPK) colouring convention for distinguishing atoms. Water molecules are not shown. **f)** Phe58:Phe126 averaged distance over unbiased molecular dynamics in the *μ*s time scale.

**Fig. 2.**
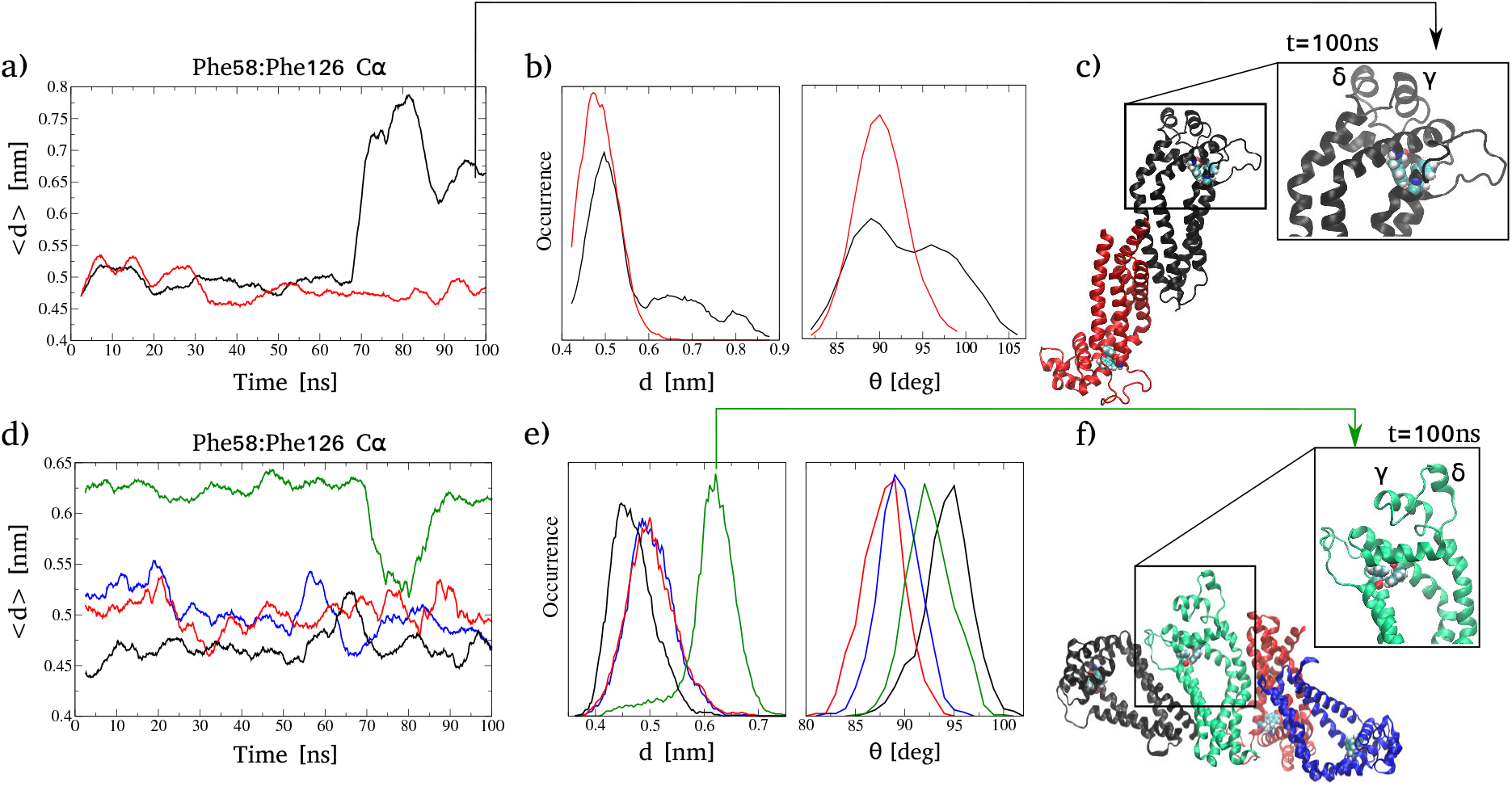
Unbiased atomistic molecular dynamics for multiple CD81 tetraspanins: Top: 2 CD81 tetraspanins. Bottom: 4 CD81 tetraspanins. **a)** and **d)** Phe58:Phe126 averaged distance. **b)** and **e)** Averaged distribution of the angle and the distance. **c)** and **f)** Molecular dynamics snapshots showing final (t=100ns) states. CD81 tetraspanins are represented in black, red, green and blue ribbons, with Phe58 and Phe126 highlighted in vdW representations following the Corey-Pauling-Koltun (CPK) colouring convention for distinguishing atoms. Water molecules are not shown.

For the system including 4 CD81 tetraspanins in water (yet, with no bilayer) the effect is even more clear. Right after minimization and equilibration (t=0ns) one of the four tetraspanins is already in its opening transition, with *d* ≈ 0.65*nm*, (panel 2d green curve). Panel 2e maps averaged distributions for *d* and *θ* of all conformations visited during these unbiased simulations, see supplementary information for 2D scattered maps. It can be observed that the cooperative interactions between four identical CD81 induce one CD81 to spontaneously increase its < *d* > distance (see green curve in the distance distribution in panel 2e) accompanied by a marginal increase in *θ* (see green curve in the angle distribution in panel 2e). Panel 2f show a significant change of conformation in the γ and δ loops, that simultaneously pull while increasing Phe58:Phe126 distance (*d*) (see supplementary information for the initial configuration).

These results are consistent with the molecular mechanism proposed by Lang and collaborators where the δ-loop flexibility is a key element to enhance CD81:CD81 direct interactions ^62^ and though to large-scale protein oligomerisation ^34,35^. Additionally, Cunha et al. showed that open-closed conformational plasticity is pH modulated ^16^ and suggested that this conformational dependence is mediated by the glycoproptein E2 in the Hepatitis C virus. Therefore, we propose that outside the lipid bilayer the opening transition is enhanced by the interplay between CD81 tetraspanins, being the event a cooperative phenomenon.

### 2.2 Segregation and opening are coupled events

To study large protein-membrane systems, coarse-grained molecular dynamics have shown to be an adequate tool, in particular, to observe emergent properties difficult to anticipate at lower scales ^46,49,63^. Here, we have performed *μ*s-length unbiased coarse-grained molecular dynamics separately for one, two, three and four CD81 embedded in a 1024 DPPC:DOPC:CHOL (0.58:0.12:0.3) lipid bilayer, solvated in explicit solvent. This ternary bilayer follows a suitable mixture used before to calculate the bending modulus in multi-component lipid membranes in different thermodynamic phases ^45^. Therefore including a saturated component with dipalmitoylphosphatidylcholine (DPPC), an unsaturated one with dioleoylphosphatidylcholine (DOPC) and (most importantly for CD81 dynamics) cholesterol (CHOL).

We have observed that CD81 tend to bind together, keeping one tetraspanin always segregated when more than 2 of them are present in a ≈ 16×16nm bilayer patch. Figure 3 shows Radial Distribution Functions (RDF) of the protein-protein distances (panels 3a,3b and 3c) and the total number of direct contacts between CD81 (panels 3d,3e and 3f), for system setups containing 2, 3 and 4 CD81 inserted in the bilayer. In all cases RDFs have been calculated between protein centres of mass (COM) and the number of CD81:CD81 direct interactions accounted only for those contacts <0.6nm. These measurements were performed over 5*μ*s of unbiased simulations.

**Fig. 3.**
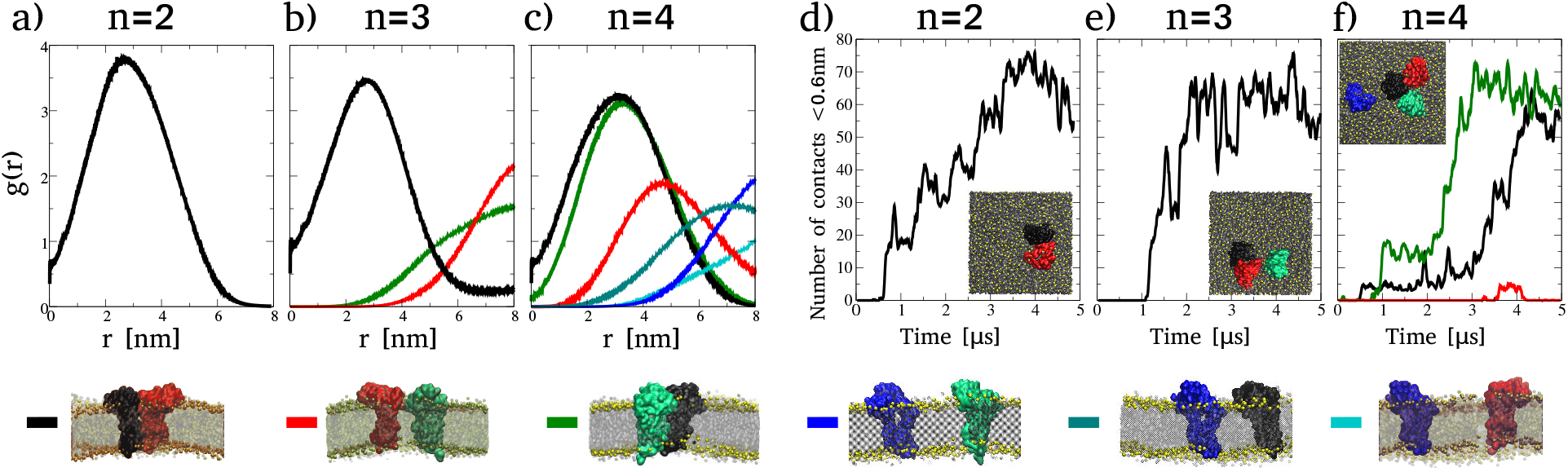
CD81:CD81 direct interactions in the lipid bilayer. Panels **a)** n=2, **b)** n=3 and **c)** n=4, are radial distribution functions of the all protein-protein distances. Measured between protein centres of mass (*r_COM_*) and calculated along 5*μ*s of unbiased simulations. Panels **d)** n=2, **e)** n=3 and **f)** n=4, are the number of direct contacts (<0.6nm) between CD81 tetraspanins. All molecular dynamics snapshots show proteins in surface representations in black, red, green and blue. The lipid bilayer is grey with PO4 beads highlighted in yellow. For clarity, water molecules are not shown.

In panel 3a the RDF of the distance between the first (black protein) and second (red protein) tetraspanin is plotted, showing its peak at ≈ 2.5nm. As expected, for n=2 both CD81 bind together exhibiting a large number of direct protein-protein contacts over time (panel 3d). In panel 3b the RDF of the distance between the third (green protein) and the first (black protein) tetraspanins has its maximum at ≈ 8nm (green curve). The same happens for the third (green protein) and the second (red protein) tetraspanin (red curve). These RDFs indicate that the third tetraspanin (green protein) is predominantly far away from tetraspanin 1 (black protein) and 2 (red protein), as confirmed by counting the number of direct contacts (panel 3e), where no green neither red curves exist. On the other hand, the black curve shows that tetraspanins 1 (black protein) and 2 (red protein) are bound together with a maximum RDF of their protein-protein COM distance ≈ 2.5nm, just like in the system with two tetraspanins inserted in the bilayer (panel 3a). Again, this result is verified by the number of direct contacts in panel 3e, (black curve).

Analogously, in panel 3c blue, cyan and turquoise curves show the RDFs of the distances measured from the fourth tetraspanin (blue protein) to the other three ones (black, red and green proteins). In all three cases the RDFs maxima are ≈ 7-8nm, indicating that the fourth tetraspanin (blue protein) remains significantly separated from the other tetraspanins during the majority of the simulation time. On the contrary, the other 3 tetraspanins have distance RDF maxima at lower values, revealing that they are permanently bound together (black, red and green curves). Panel 3f confirms these RDF results by measuring <0.6nm CD81:CD81 contacts for n=4. In the snapshot inset it is observed that 3 tetraspanins form an assembly (black, red and green proteins) leaving one tetraspanin (blue protein) out of their group. Accordingly, no contacts <0.6nm between the blue tetraspanin and any other one exist (no blue, nor cyan, nor turquoise curves in panel 3f). Noticeably, in this ternary assembly one of the tetraspanins acts as a bridge-linker (black protein) binding together the other two (red and green proteins), as indicated by the small amount of direct contacts between these two tetraspanins (red curve in panel 3f).

Remarkably, the tetraspanin that is segregated in systems with n>2 appears to more readily adopt an open conformation than the ones bound together. Figure 4 shows Phe58:Phe126 distances for each CD81 in the same 4 simulation setups: the lipid bi layer containing either 1, 2, 3 or 4 identical CD81. Panels 4c and 4d show one CD81 with significantly larger Phe58:Phe126 distance maintained during 5*μ*s of unbiased simulations. Panel 4c shows Phe58:Phe126 distance (green curve) during the segregation of the third tetraspanin (green protein) and panel 4d the Phe58:Phe126 distance (blue curve) during the segregation of fourth tetraspanin (blue protein). These two tetraspanins (green and blue proteins) are the same ones that systematically locate apart from the others in previous figure 3 for n=3 and n=4.

**Fig. 4.**
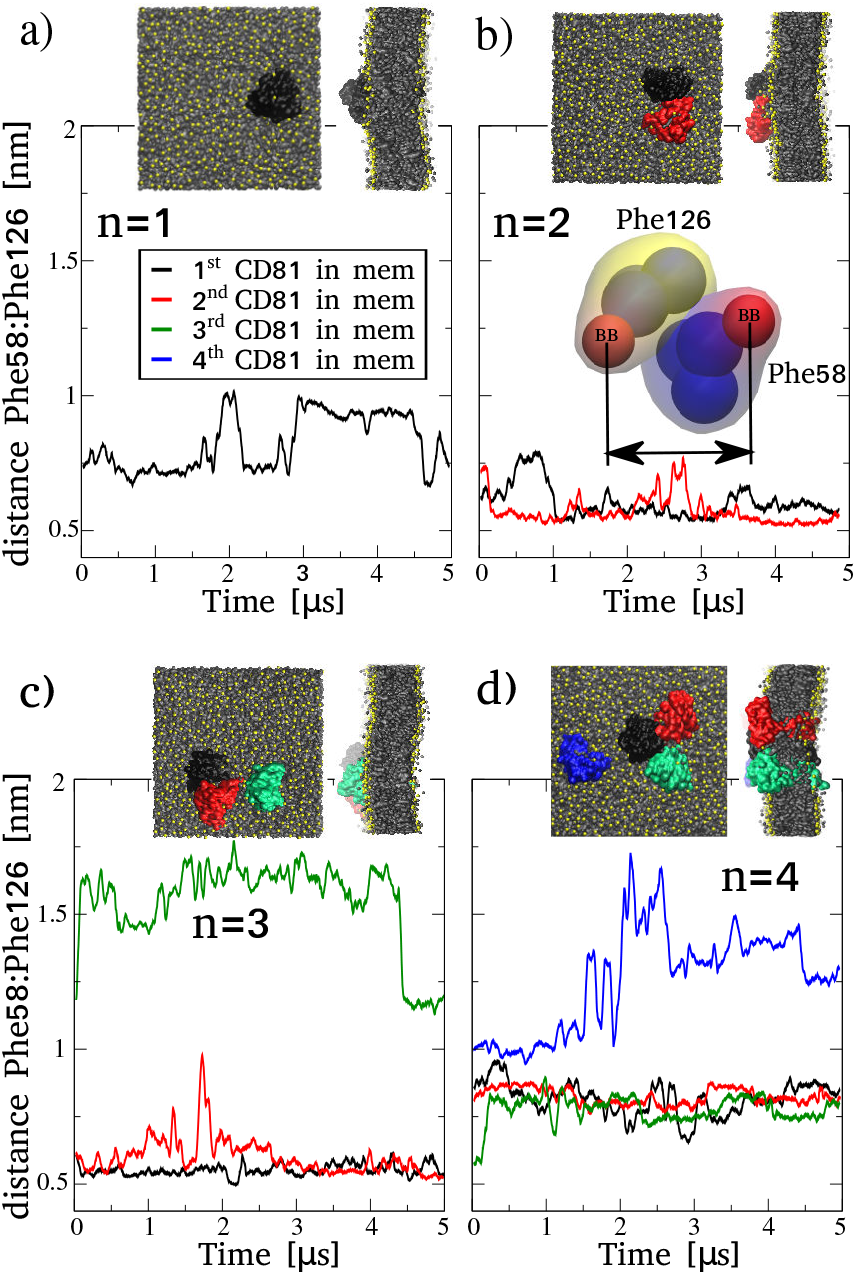
Phe58:Phe126 distances for each individual CD81 embedded in the membrane. Measured between BB beads in Phe58 and Phe126 residues (see inset in panel b) along 5*μ*s of unbiased simulations for 4 different systems with increasing amounts of CD81: **a)** n=1, **b)** n=2, **c)** n=3 and **d**) n=4. All molecular dynamics snapshots show proteins in surface representations in black, red, green and blue. The lipid bilayer is grey with PO4 beads highlighted in yellow. Top and side views are provided. For clarity, water molecules are not shown.

In summary, we have observed again a cooperative phenomenon, this time for CD81 inserted in a ternary bilayer including cholesterol. Although proteins tend to cluster in all cases, when n>2 a single tetraspanin is regularly left apart. Notably, that segregated tetraspanin exhibits a significant increase in its Phe58:Phe126 distance showing a first-stage or partial-switch to an open conformation. Such collective protein-protein partnership is again in line with the capabilities of tetraspanins to oligomerize via the flexibility of the external loops ^62^, forming tetraspanin webs ^35^ and organizing themselves in homomeric clusters ^34^.

### 2.3 CD81 sequesters membrane cholesterol and inhibits large deformations

Like other members of the tetraspanin superfamily (*i.e.* CD9^69^), it has been suggested that CD81 could influence the curvature of the cell membrane ^17,23,75^. Here, to induce bilayer curvature and to quantify its associated free-energy cost, we have used a collective variable specifically design for that scope, which we have already used in other proteins adsorbed on the membrane surface ^13,51^. The collective variable, defined **Ψ**, forces lipid molecules to increase their local density at the centre of the bilayer, inducing the membrane to make slight displacements along its normal axis (Z). This rearrangement of lipids bends the membrane in a saddle-like shape which emerges spontaneously ^18,52^. Unlike other approaches ^72^, **Ψ** does not allow for a direct control of the bilayer curvature but is rather an indirect mechanism analogous to the protein-membrane surface interplay, as previously described for α-synuclein ^13,36^ and the N-BAR domain ^52^, both known for inducing curvature in lipid bilayers ^57,74^. Besides, the complex saddle-shape mode in which the bilayer bends follows two principal curvatures defined by two fitted circles with different radii ^3,4,22^ and requires significantly more energy than other modes of simpler geometry.

Mathematically, **Ψ** is a dimensionless reaction coordinate defined by eq. 1:

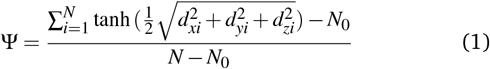

Here, the sum ranges over the centres of mass of every lipid molecule in the bilayer (*N* = 1024). Distances *d_xi_*, *d_yi_* and *d_zi_* are the XYZ components of the three-dimensional distance between the centre of mass of bilayer and the centre of mass of lipid molecule *i*. Constant *N*_0_ is the average equilibrium value of the summation in an unbiased simulation of a flat bilayer (for this bilayer, *N*_0_=991.771). For technical details on the development and implementation of **Ψ**, see Masone et al. ^52^.

By exploring the phase space in **Ψ**, we have found that a single CD81 tetraspanin increases the free energy cost to bend the bilayer in ≈ 200 *k_B_T* (see figure 5, black curve) with respect to the same value of **Ψ** ≈ 0.49 in an identical bilayer-only system (magenta curve). Given that **Ψ** is a function of the membrane (not the protein) any changes in the free energy are exclusively due to the interference of CD81 in the bilayer dynamics.

**Fig. 5.**
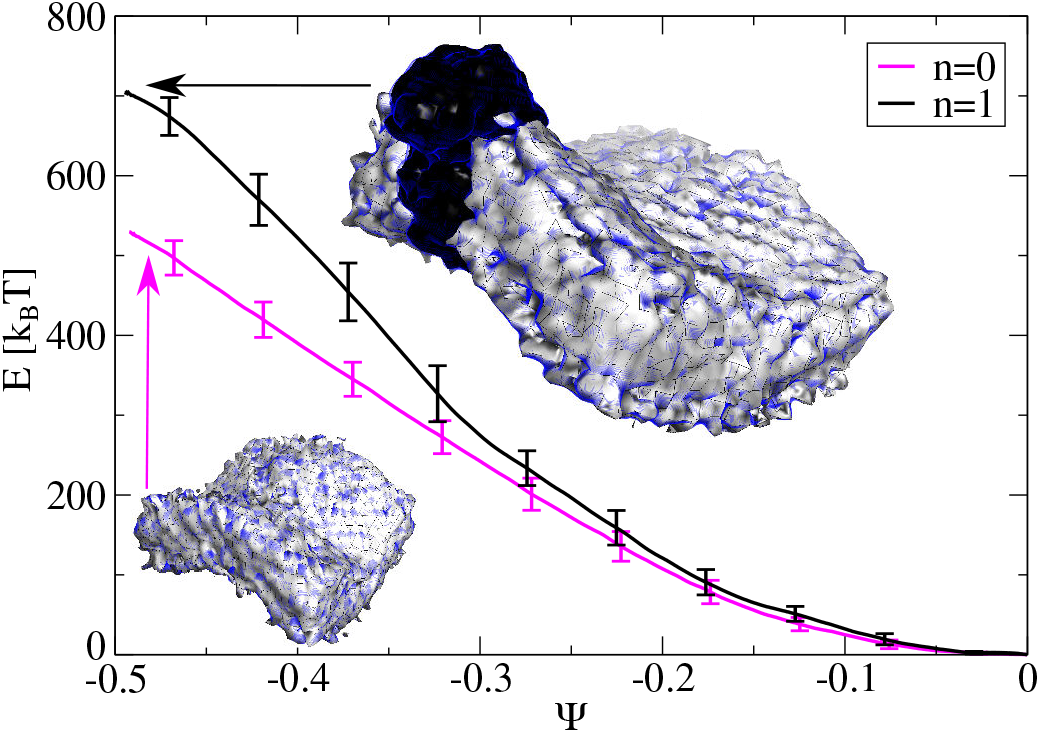
PMF calculations for membrane bending in the Ψ space. Membrane-only (magenta, n=0) and membrane with a single CD81 (black, n=1). Error bars are standard errors calculated by individually splitting the profiles in independent blocks. Insets show the bilayer with (top) and without (bottom) CD81. Snapshots are averaged densities for all lipid molecules showing steady-state curvature for Ψ = −0.49. See supplementary information for PMF convergence details.

The reason for this free-energy increment during bending is explained by the strong interactions taking place between CD81 transmembrane region and the surrounding cholesterol molecules. One single tetraspanin transmembrane domain captures between 11 and 13 cholesterol molecules that remain strongly attached to the protein surface (see panel 6b) and are unable to move along the normal axis to the membrane, though resisting to the curvature mechanism imposed by the reaction coordinate. Moreover, at least 3 of these cholesterol molecules remain trapped inside the protein pocket (see detail in panel 6b).

To quantitatively demonstrate this effect, we have calculated the Mean Square Displacement (MSD) for different groups of cholesterol molecules (all cholesterol molecules in the bilayer and cholesterol molecules inside the protein binding pockets, see panel 6a) to estimate their degree of adherence to CD81 in unbiased 5*μ*slength simulations. Then, the cholesterol lateral diffusion coefficient in [nm^2^/ms] can be extracted by fitting a straight line to the MSD at the linear region, in other words, the lateral diffusion coefficient is the slope of the linear region of MSD plot. Comparing MSDs in panel 6a, it is observed that cholesterol molecules inside the protein binding pockets exhibit much lower values (red curve), though a much lower lateral diffusion coefficient is expected when compared to the complete set of cholesterol molecules in the bilayer (black curve), quantitatively verifying the confinement phenomenon.

For more CD81 tetraspanins in the membrane the bending-resistant effect increases drastically. As more cholesterol molecules strongly bind to more transmembrane domains, more difficult is for the membrane to bend as a response to the local density increase imposed by **Ψ**. Without increasing the force constant used by the reaction coordinate, with two, three and four CD81 the bilayer is unable to bend. Figure 7 shows the lipid bilayer with n=1, 2, 3 and 4 CD81 under the same harmonic restrain applied with **Ψ**. The saddle-shaped curvature emerges only for n=1 and there is no significant permanent curvature for n≥2. As expected, the density control strategy used by **Ψ** to successfully induce curvature in membrane-only systems and in membranes with surface proteins ^13,52^, is almost nullified here by the strong protein-cholesterol binding and further confinement inside the binding pockets. This is observed for more than one CD81 inserted in this ≈16×16nm bilayer patch containing 306 cholesterol molecules (see figure 7).

To evaluate cholesterol molecules fluctuations in 2D, figure 8 shows density maps for the Root Mean Square Fluctuations (RMSF) of all cholesterol molecules in the bilayer, averaged along 5*μ*s of unbiased molecular dynamics, for 4 setups: the bilayer with 1, 2, 3 or 4 CD81 tetraspanins. In all cases, less fluctuations are observed for molecules located near the transmembrane protein domains (violet regions). This result describes the unique mechanism used by CD81 to interact with the bilayer. Not only an amount of cholesterol molecules are slowed down in the surroundings of each protein, but also each CD81 sequesters a small amount of cholesterol inside its biding pocket (see figure 6). This is consistent with other reports highlighting the strong link between cholesterol and tetraspanin biology ^15,19,64^.

**Fig. 6.**
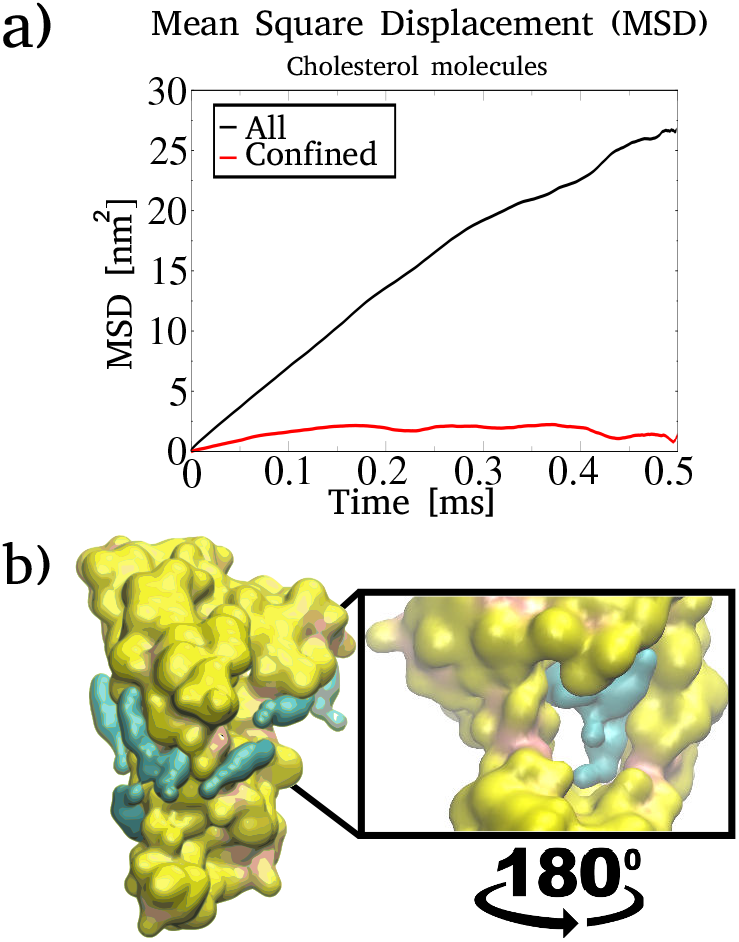
Transmembrane region in CD81 captures cholesterol molecules. **a)** Mean square displacement of all cholesterol molecules in the bilayer (black) and cholesterol molecules inside the protein pocket (red). **b)** One CD81 (yellow) with cholesterol molecules (cyan) bound to its transmembrane domain and the detail of cholesterol molecules sequestered inside the protein pocket.

**Fig. 7.**
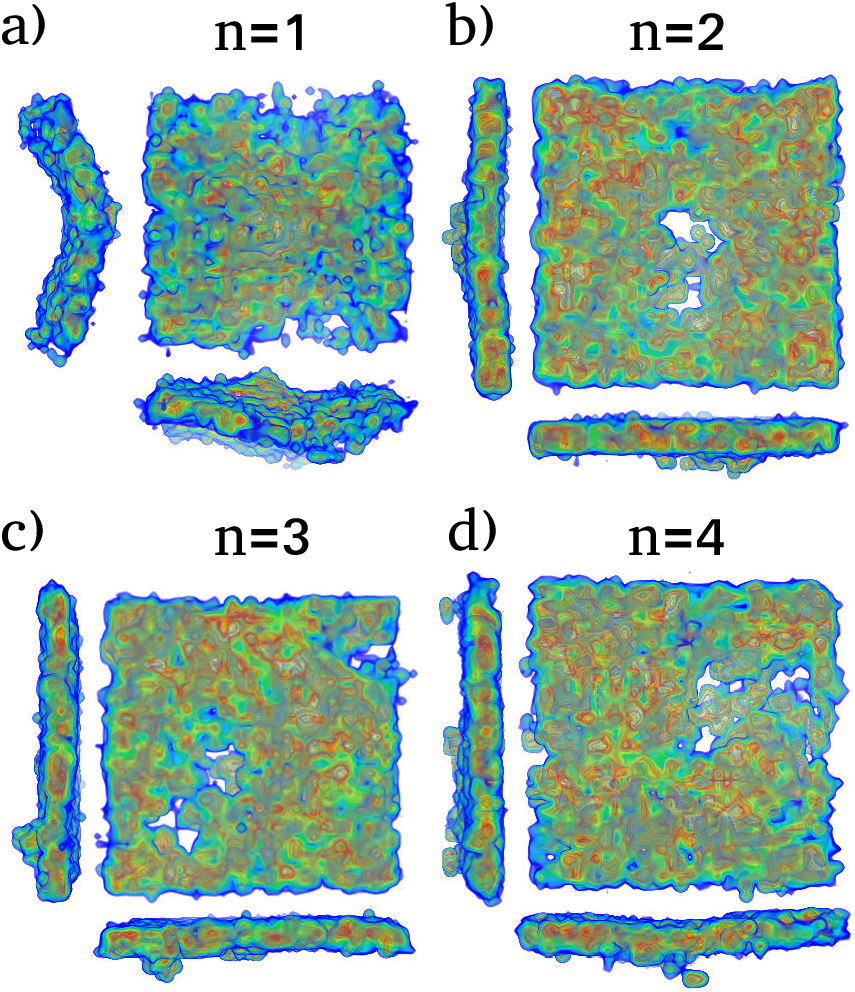
Four membranes with increasing amounts of CD81 (n=1, 2, 3 and 4) under identical restrain conditions. (Ψ = −0.49). Averaged cholesterol-only density maps depicting how the membrane is unable to bend in its characteristic saddleshape, when 2 or more CD81 are inserted in the bilayer (n≥2).

**Fig. 8.**
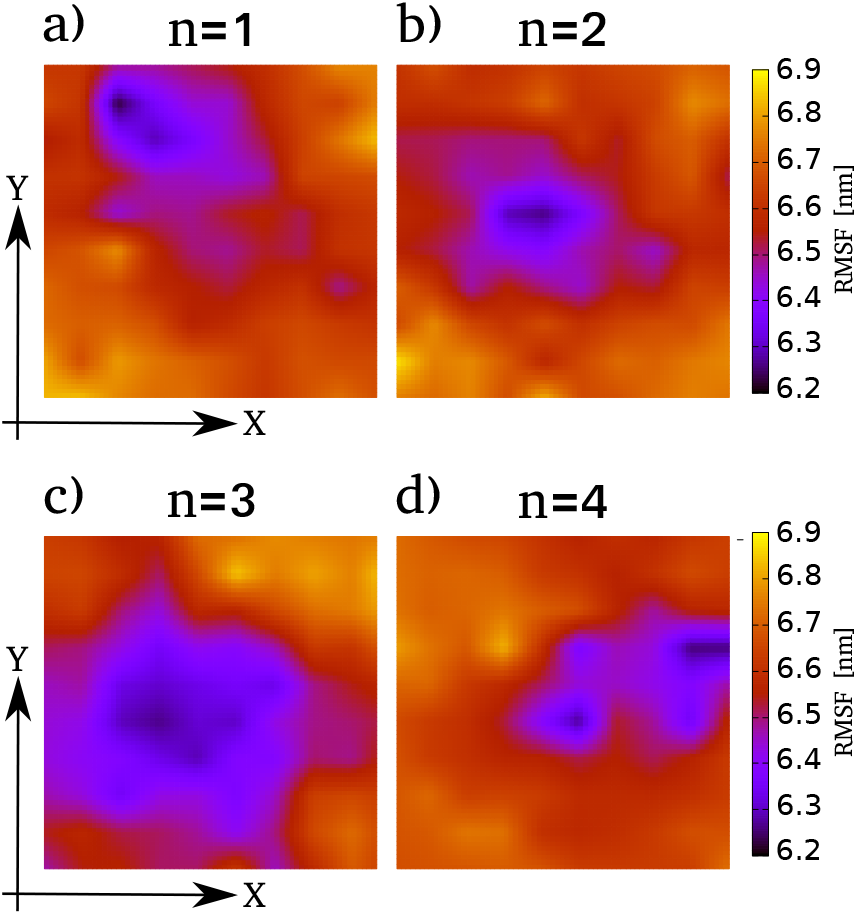
Root Mean Square Fluctuations (RMSF) density maps for all cholesterol molecules. Averaged along 5*μ*s of unbiased simulations for 4 different systems with increasing amounts of CD81: **a)** n=1, **b)** n=2, **c)** n=3 and **d**) n=4.

## 3 Conclusions

In this work we have studied the collective properties of CD81 tetraspanin clusters, both with and without the bilayer. We have shown that the conformational opening transition of CD81, known to be important in protein oligomerisation, is facilitated by CD81 clusters as a collaborative phenomenon, less likely to occur for an isolated CD81 outside the bilayer. We have observed that assemblies of 2 and 4 identical tetraspanins form groups with higher propensity to induce one tetraspanin into an initial-stage opening transition, still with no membrane interference.

Simulations of multiple CD81 inserted in a ternary DPPC:DOPC:CHOL (0.58:0.12:0.3) bilayer verified the clustering behaviour observed before. Interestingly, for more than two identical tetraspanins inserted in the membrane, CD81 assemblies tend to segregate one single tetraspanin that again shows an advanced opening transition, an emergent property not observed individually. Finally, by inducing membrane curvature we have described at molecular level the strong protein-cholesterol interactions that lead to cholesterol sequestering by CD81, inhibiting large membrane deformations due to local density changes.

In conclusion, our study shows that CD81 tetraspanin dynamics are highly dependent on their mutual interactions and their interplay with the cholesterol-containing lipid bilayer. We suggest that emergent properties arising from complex group dynamics are key in these proteins inclined to form enriched domains.

## 4 Computational methods

All simulations were performed with GROMACS-2018.3^1,58,70^ patched with Plumed 2.4.3^68^, a plug-in for free energy calculations and complex collective variable implementation. The calculation of **Ψ** was implemented in Plumed, which by using Lepton C++ Mathematical Expression Parser is able to evaluate symbolic expressions, such as the hyperbolic tangent used by **Ψ** as switching function. Radial distribution functions were calculated with GRO-MACS built-in *gmx rd f* while inter-residue distances were measured with *gmx mindist*. RMSF calculations were performed using *gmx rms f*. Coarse-grained bilayers and proteins were prepared in part using CHARMM-GUI web server ^43^. Molecular dynamics snapshots were visualized using Visual Molecular Dynamics (VMD) ^38^. Data graphics were plotted with Grace (GRaphing, Advanced Computation and Exploration of data). Figure panels were organized using Inkscape. Density maps from molecular dynamics simulations were generated using Gromaρs ^9^.

### 4.1 Atomistic molecular dynamics

All atomistic molecular dynamics were performed in the isotropic NPT ensemble at T = 303.15K ^42,44,61^ using the Nosé-Hoover thermostat ^53^ with a relaxation time of 1ps. The compressibility was set to 4.5×10^−5^ bar^−1^ using a Parrinello-Rahman barostat ^56^ with a time constant of 5ps and a reference pressure of 1 bar. Hydrogen bond lengths were constrained using the LINCS algorithm. Lennard-Jones and real-space electrostatic interactions radii were set to 1.2nm. Hydrogen bond lengths were constrained using the LINCS algorithm ^32^. The Verlet cutoff scheme ^54^ was used updating the neighbour lists every 20 steps. Particle Mesh Ewald (PME) method ^20^ was used for long-range electrostatic interactions. Periodic boundary conditions were applied in all three dimensions. The Transferable Intermolecular Potential with 3 points (TIP3) model was used for water molecules. The chosen force field in all cases was CHARMM36^10^. For simulations including 1 CD81 tetraspanin solvated in ≈45000 water molecules the simulation box length resulted in L≈11nm (cubic shape). For simulations including 2 CD81 tetraspanin solvated in ≈51000 water molecules the simulation box length resulted in L≈12nm. For simulations including 4 CD81 tetraspanin solvated in ≈33000 water molecules the simulation box length resulted in L≈22nm. Values were set by the CHARMM-GUI protein solvation algorithm ^41^. Unbiased atomistic simulations including 1 CD81 tetraspanin solvated in explicit TIP3 water run for 1 us. Umbrella windows for the free energy profile in figure 1 run for 100ns each, adding a total of 1.5us. For 2 and 4 CD81 tetraspanin solvated in explicit TIP3P water, unbiased simulations run for 100ns. All atomistic production runs used a time step of 2fs and collected data every 50000 steps.

### 4.2 Coarse-grained molecular dynamics

The MARTINI coarse-grained model ^50^ is widely used for protein-lipid molecular modelling ^14,26,31,40,71^. In this particular reduced resolution space, beads are classified according to their mutual interactions as: polar, non-polar, apolar and charged. Cholesterol molecules are represented by 8 beads (6 modelling the sterol and 2 the tail) while keeping a mainly planar geometry. Phosphatidylcholine (PC) headgroups include two hydrophilic groups: choline charged positively and phosphate charged negatively ^50^.

All production molecular dynamics simulations were run in the semi-isotropic NPT ensemble at T=303.15K with the V-rescale thermostat ^11^ and a relaxation time of 1ps. The compressibility was set to 3×10^−4^ bar ^−1^, using a Parrinello-Rahman barostat ^56^ with a time constant of 12ps. Long-range reaction field electrostatic interactions were used with a Coulomb cut-off set to 1.1nm. Before production runs, all systems were minimized and equilibrated for at least 100ns to generate properly relaxed initial configurations. Periodic boundary conditions were applied in all three dimensions. For equilibration runs the pressure was set at 1.0 bar using the Berendsen barostat ^5^ with a 5ps coupling constant. In all cases we used the polarisable water model for MARTINI ^73^. All bilayers containing 1024 molecules were solvated in ≈ 18×10^3^ water molecules, satisfying the ample water condition for MARTINI ^39^. In the coarse-grained space the simulation box dimensions resulted in Lx≈16nm, Ly≈16nm and Lz≈12nm. All unbiased simulations in the coarse-grained space run for 5us each. Umbrella windows for the free energy profile in figure 5 run for 100ns, adding a total of 5.1us. As suggested for MARTINI, all coarse-grained production runs used a time step of 20fs and collected data every 5000 steps.

Due to particular artifacts pointed out in the MARTINI model regarding the overestimation of protein-protein and protein-membrane interactions ^2,40^, we have back-converted final configurations (t=5*μ*s) to atomistic resolution using the CHARMM-GUI all-atom converter ^41^ for MARTINI. We have applied this conversion to systems containing 1, 2, and 4 CD81 tetraspanins inserted in the DPPC:DOPC:CHOL bilayer fully solvated in water, following the same arrangement used along the atomistic simulations to verify their physical feasibility. Once in the atomistic space, all configurations were relaxed with a short minimization using the steepest descend method and subsequently equilibrated under NVT and NPT conditions with time steps of 1fs and 2fs. Plots of the area of the atomistic membranes show proper relaxation in all cases (see supplementary information). No protein-protein unbinding nor protein-membrane artifacts were observed. Molecular dynamics snapshots for these simulations are provided in supplementary information.

### 4.3 PMF calculations

PMF profiles were computed with umbrella sampling ^60,67^ and were recovered using the Weighted Histogram Analysis Method (WHAM) ^48^ in GROMACS as *gmx wham* ^37^ and also with the WHAM implementation by Prof. Grossfield version 2.0.9.1^24^. Convergence was assessed by repeating these calculations on consecutive trajectory blocks. For technical details on umbrella sampling windows distribution and force constants values see supplementary information.

## Acknowledgements

Supercomputing time for this work was provided by CCAD (Centro de Computación de Alto Desempeño de la Universidad Nacional de Córdoba) and by SNCAD-MinCyT Iniciativa de Proyectos Acelerados de Cálculo (IPAC), grant numbers: 2017-75-APN-SECACT#MCT, 2018-6-APN-SECACT#MECCYT and 2019-72-APN-SECACT#MECCYT. Also, grants from CONICET (PIP-13CO01), ANPCyT (PICT2017-1002) and GPU hardware by the NVIDIA Corporation are gratefully acknowledged.

