## Supplementary material for "Cholesterol plays a decisive role in tetraspanin assemblies during bilayer deformations"

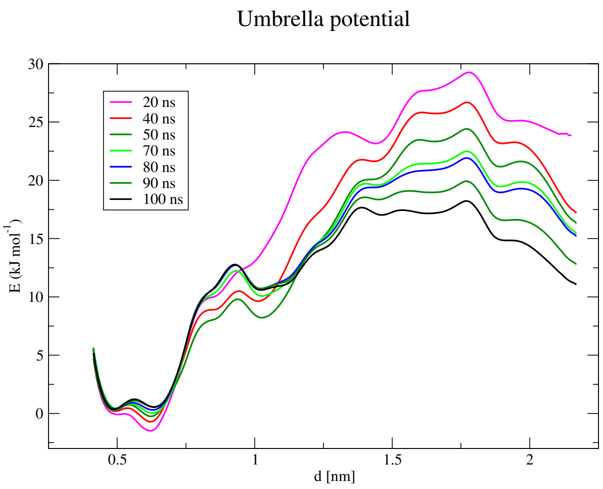


Figure S1: **Convergence of the free energy profile for CD81 opening transition.** Calculated with *gmx wham* over increasing lengths of simulation time.

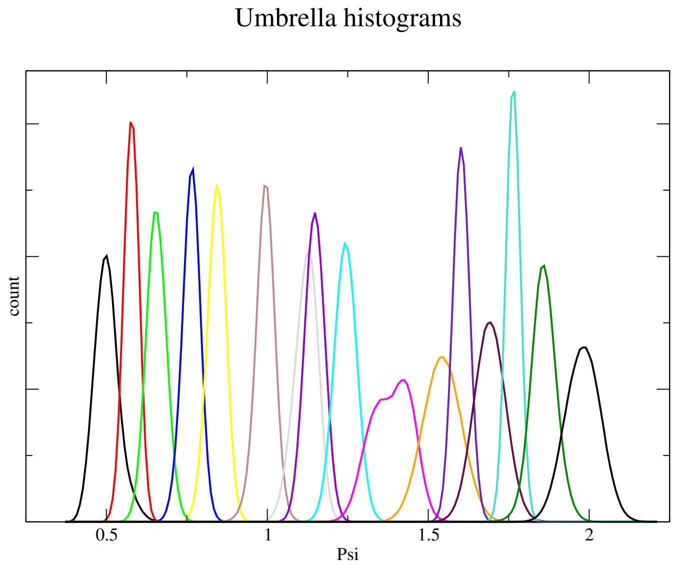


Figure S2: **Histogram distribution.** Reaction coordinate $d$ spans from 0 to 2nm in 16 umbrella windows.

Table S1: Force constants and equilibrium restrains for PMF in CD81 opening transition

| window | $k[kJ/mol/nm^2]$ | $d_0$ |
|---|---|---|
| 0 | 1000 | 0.5 |
| 1 | 5000 | 0.6 |
| 2 | 2000 | 0.7 |
| 3 | 3000 | 0.8 |
| 4 | 2000 | 0.9 |
| 5 | 2000 | 1.0 |
| 6 | 2000 | 1.1 |
| 7 | 3000 | 1.2 |
| 8 | 2000 | 1.3 |
| 9 | 1000 | 1.4 |
| 10 | 1000 | 1.5 |
| 11 | 4000 | 1.6 |
| 12 | 1000 | 1.7 |
| 13 | 5000 | 1.8 |
| 14 | 1000 | 1.9 |
| 15 | 1000 | 2.0 |

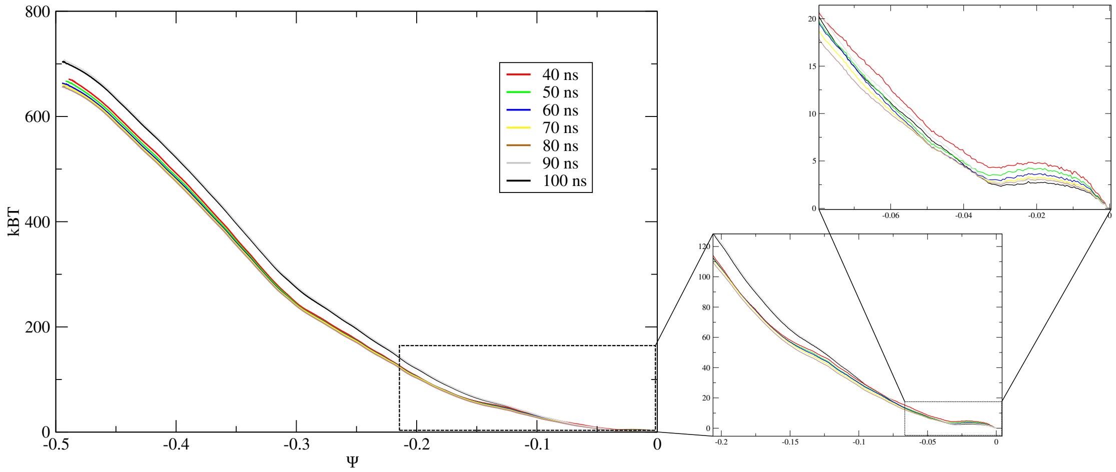


Figure S3: **Convergence of the free energy profile for bending the lipid bilayer.** Calculated with Prof. Grossfield's WHAM implementation over increasing lengths of simulation time.

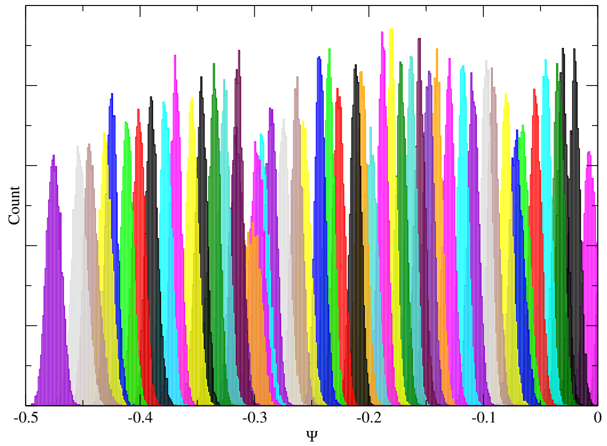


Figure S4: **Histogram distribution.** Reaction coordinate Ψ spans from -0.5 to 0 in 51 equidistant umbrella windows, in steps of 0.01 using in all cases a force constant k=100000 [kJ/mol].

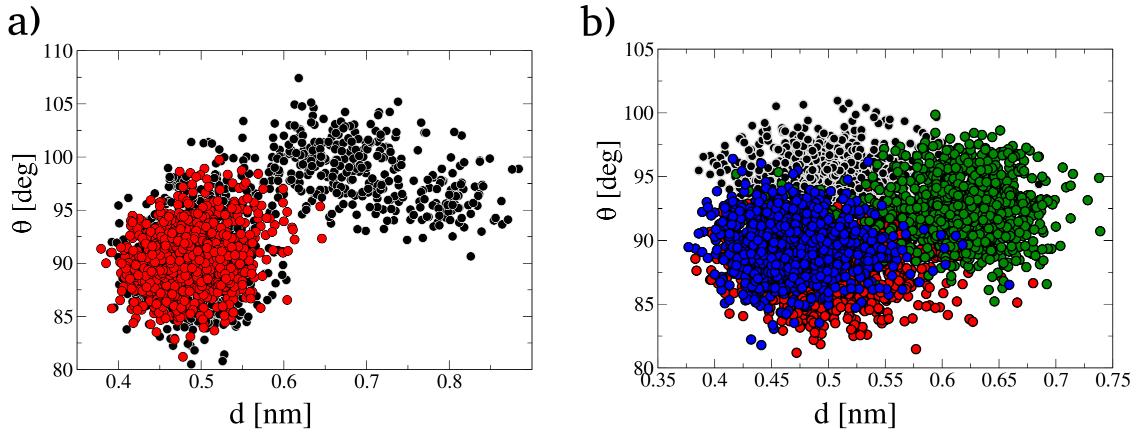


Figure S5: **Distance-angle 2D scattered maps for atomistic simulation.** **a)** 2 CD81 tetraspanins. **b)** 4 CD81 tetraspanins. In both cases colors match protein colors in manuscript figure 2.

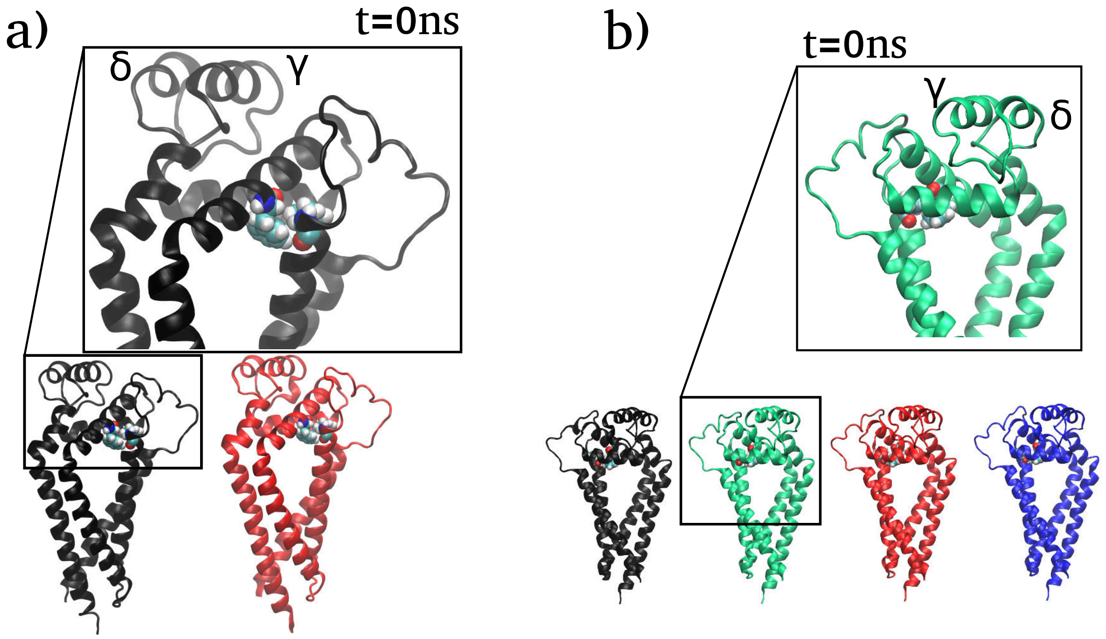


Figure S6: **Molecular dynamics snapshots showing initial (t=0ns) states.** **a)** 2 CD81 tetraspanins. **b)** 4 CD81 tetraspanins. CD81 tetraspanins are represented in black, red, green and blue ribbons, with Phe58 and Phe126 highlighted in vdW representations following the Corey-Pauling-Koltun (CPK) colouring convention for distinguishing atoms. Water molecules are not shown.

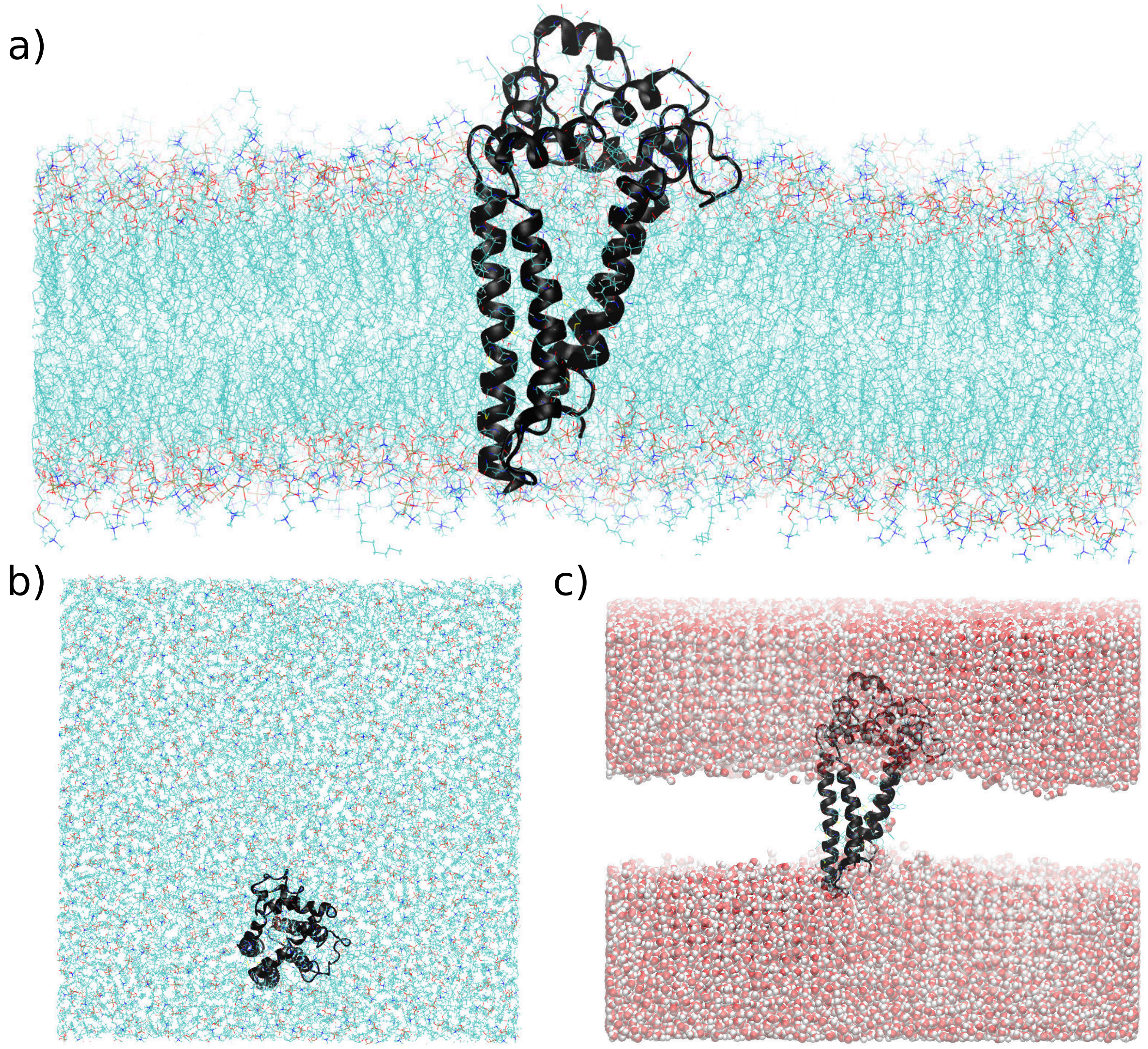


Figure S7: **Atomistic molecular dynamics snapshots back-converted from the coarse-grained space for 1 CD81 inserted in the bilayer. a)** Side-view of the membrane with the protein (black ribbons). **b)** Top view. Lipid molecules are in line representation with Corey-Pauling-Koltun (CPK) colouring convention for distinguishing atoms. **c)** Side-view of water molecules (in vdW representation) with the protein.

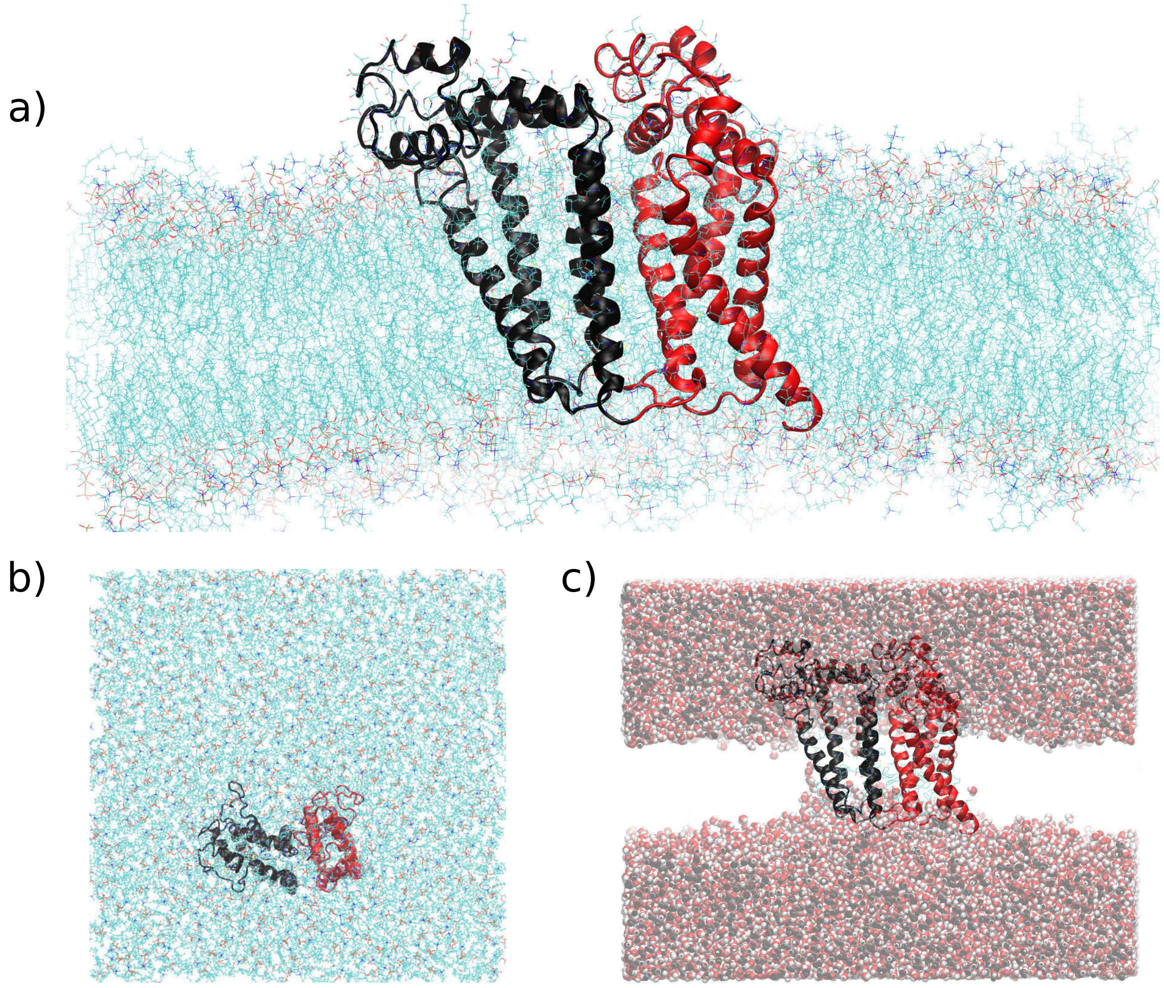


Figure S8: **Atomistic molecular dynamics snapshots back-converted from the coarse-grained space for 2 CD81 inserted in the bilayer. a)** Side-view of the membrane with the protein (black and red ribbons). **b)** Top view. Lipid molecules are in line representation with Corey-Pauling-Koltun (CPK) colouring convention for distinguishing atoms. **c)** Side-view of water molecules (in vdW representation) with the proteins.

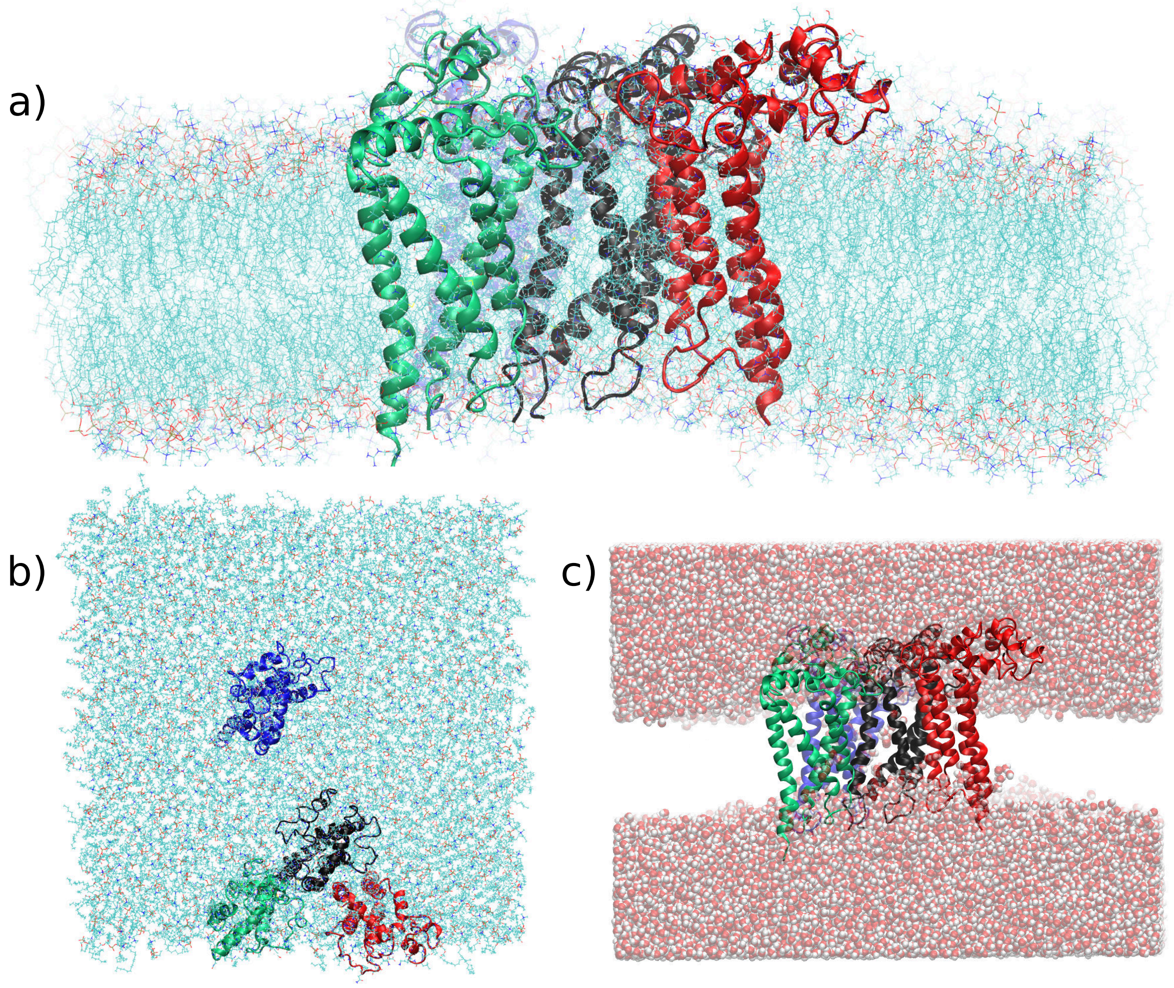


Figure S9: **Atomistic molecular dynamics snapshots back-converted from the coarse-grained space for 4 CD81 inserted in the bilayer. a)** Side-view of the membrane with the protein (black, red, green and blue ribbons). **b)** Top view. Lipid molecules are in line representation with Corey-Pauling-Koltun (CPK) colouring convention for distinguishing atoms. **c)** Side-view of water molecules (in vdW representation) with the proteins.

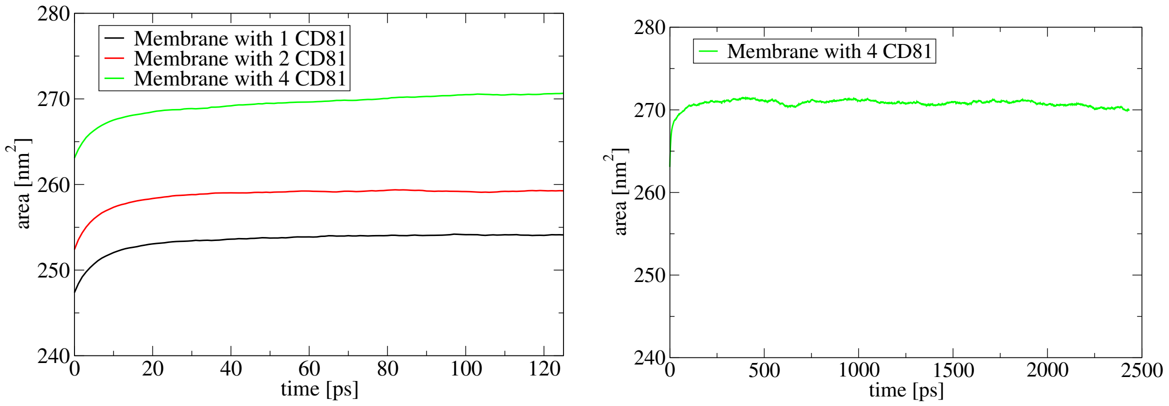


Figure S10: **Area fluctuations in the atomistic bilayers. Left:** Short equilibration for area relaxing in 3 protein-membrane systems. **Right:** Extension by 20 times to verify equilibrated membrane area for the system with the maximum quantity of proteins inserted.
